# Adaptive laboratory evolution restores solvent tolerance in plasmid-cured *Pseudomonas putida* S12; a molecular analysis

**DOI:** 10.1101/2020.08.01.232264

**Authors:** Hadiastri Kusumawardhani, Benjamin Furtwängler, Matthijs Blommestijn, Adelė Kaltenytė, Jaap van der Poel, Jevan Kolk, Rohola Hosseini, Johannes H. de Winde

## Abstract

*Pseudomonas putida* S12 is intrinsically solvent-tolerant and constitutes a promising platform for biobased production of aromatic compounds and biopolymers. The genome of *P. putida* S12 consists of a 5.8 Mbp chromosome, and a 580 kbp megaplasmid pTTS12 that carries several gene clusters involved in solvent tolerance. Removal of pTTS12 caused a significant reduction in solvent tolerance. In this study, we succeeded in restoring solvent tolerance in plasmid-cured *P. putida* S12 using adaptive laboratory evolution (ALE), underscoring the innate solvent-tolerance of this strain.

Whole genome sequencing revealed several single nucleotide polymorphisms (SNPs) and a mobile element insertion, enabling ALE-derived strains to survive and sustain growth in the presence of a high toluene concentration (10% v/v). Mutations were identified in an RND efflux pump regulator *arpR*, resulting in constitutive upregulation of the multifunctional efflux pump ArpABC. SNPs were also found in the intergenic region and subunits of ATP synthase, RNA polymerase subunit β’, global two-component regulatory system (GacA/GacS) and a putative AraC-family transcriptional regulator Afr. RNA-seq analysis further revealed a constitutive down-regulation of energy consuming activities in ALE-derived strains, including flagellar assembly, F0F1 ATP synthase, and membrane transport proteins. Out results indicate that constitutive expression of an alternative solvent extrusion pump in combination with high metabolic flexibility ensures restoration of solvent-tolerance in *P. putida* S12 lacking its megaplasmid.

## Introduction

*Pseudomonas putida* is a promising microbial host for biobased production of valuable chemicals and biopolymer compounds [1]. Endowed with its natural versatility, *P. putida* is robust towards toxic compounds which may arise in whole-cell biocatalysis processes as substrates, intermediates, or products. *P. putida* displays a remarkable intrinsic oxidative stress- and solvent-tolerance. This may be further optimized for utilization of secondary feedstock as carbon source and production of various aromatic compounds and bioplastics monomers [2–9]. Moreover, several metabolic models and genetic tools are currently available for the design and implementation of novel biosynthetic pathways in *P. putida* [10–12].

*P. putida* S12 was isolated from soil on minimal media to use styrene as its sole carbon source [13]. It is highly tolerant towards organic solvents and aromatic compounds which are often toxic towards microbial hosts. As such, this strain has been used to produce a variety of high value aromatic compounds [6, 8, 9, 14]. Organic solvents and aromatic compounds are toxic to most bacteria as these compounds are able to accumulate in the bacterial membrane and thus alter membrane integrity [15], resulting in damage and loss of various membrane functions such as permeability barrier, matrix for protein and metabolic reaction, energy transduction and denaturation of essential enzymes.

The *P. putida* S12 genome comprises of a 5.8 Mbp chromosome and a single-copy 583 kbp megaplasmid pTTS12 [16]. Plasmid pTTS12 encodes, among others, an RND efflux pump (SrpABC), a styrene–phenylacetate degradation pathway, and a toxin-antitoxin module slvTA are responsible for high solvent tolerance of *P. putida* S12 [13, 17, 18]. A significant reduction of solvent-tolerance was previously demonstrated when *P. putida* S12 was cured from its megaplasmid. As a result of plasmid-curing, *P. putida* S12 ΔpTTS12 could only survive and sustain growth in a maximum of 0.15% v/v toluene [18]. As a comparison, wild-type S12 can sustain growth in 0.30% toluene and survive up to 10% toluene. However, as was previously demonstrated, the expression of SrpABC efflux pump in *E. coli* and the non-solvent-tolerant *P. putida* KT2440 instigated still a lower solvent-tolerance than in *P. putida* S12, indicating the intrinsic solvent-tolerance of *P. putida* S12 [18].

In this paper, we further addressed the intrinsic solvent tolerance of *P. putida* S12. Megaplasmid pTTS12 may confer genetic adaptation towards environmental chemical stressors like organic solvents and aromatic compounds, through horizontal gene transfer. Here, we examined the ability of plasmid-cured *P. putida* S12 to survive and sustain growth in the presence of toluene. Using adaptive laboratory evolution (ALE), we were able to restore the solvent tolerance in *P. putida* S12 lacking the megaplasmid. Specific mutations putatively responsible for the restored solvent-tolerance trait were characterized. Moreover, RNA-seq transcriptional analysis revealed the constitutive responses of the plasmid-cured *P. putida* S12 after adaptation to the elevated toluene concentration.

## Materials and Methods

### Strains and culture conditions

Strains and plasmids used in this paper are listed in Table 1. *P. putida* strains were grown in Lysogeny Broth (LB) on 30 °C with 200 rpm shaking. *E. coli* strains were cultivated in LB on 37 °C with 250 rpm. For solid cultivation, 1.5 % (w/v) agar was added to LB. When required, gentamycin (25 mg l^−1^), ampicillin (100 mg l^−1^), kanamycin (50 mg l^−1^), and streptomycin (50 mg l^−1^) were added to the media. Hartman’s minimal medium [13] was supplemented with 2 mg MgSO_4_ and 0.2 % w/v of citrate, 0.4% w/v of glycerol, or 0.2% w/v of glucose may be added as sole carbon source. Growth parameters were measured in a 96-well plates using a Tecan Spark 10M instrument and calculated using growthcurver R-package ver.0.3.0 [19]. Maximum growth rate (μmax) was calculated as the fastest growth rate when there were no restrictions imposed on total population size (t = 2-5 hours). Maximum OD_600_ (maxOD) was defined as the OD_600_ measurement after the stationary phase was reached (t ≈ 10 hours). Solvent tolerance analysis was performed by growing 20 ml of *P. putida* culture (starting OD_600_ ± 0.1) on LB with the addition of toluene (0.15 – 10% v/v) in 250 ml Boston glass bottles with Mininert^®^ valve (Sigma-Aldrich) bottle caps.

**Table 1.**
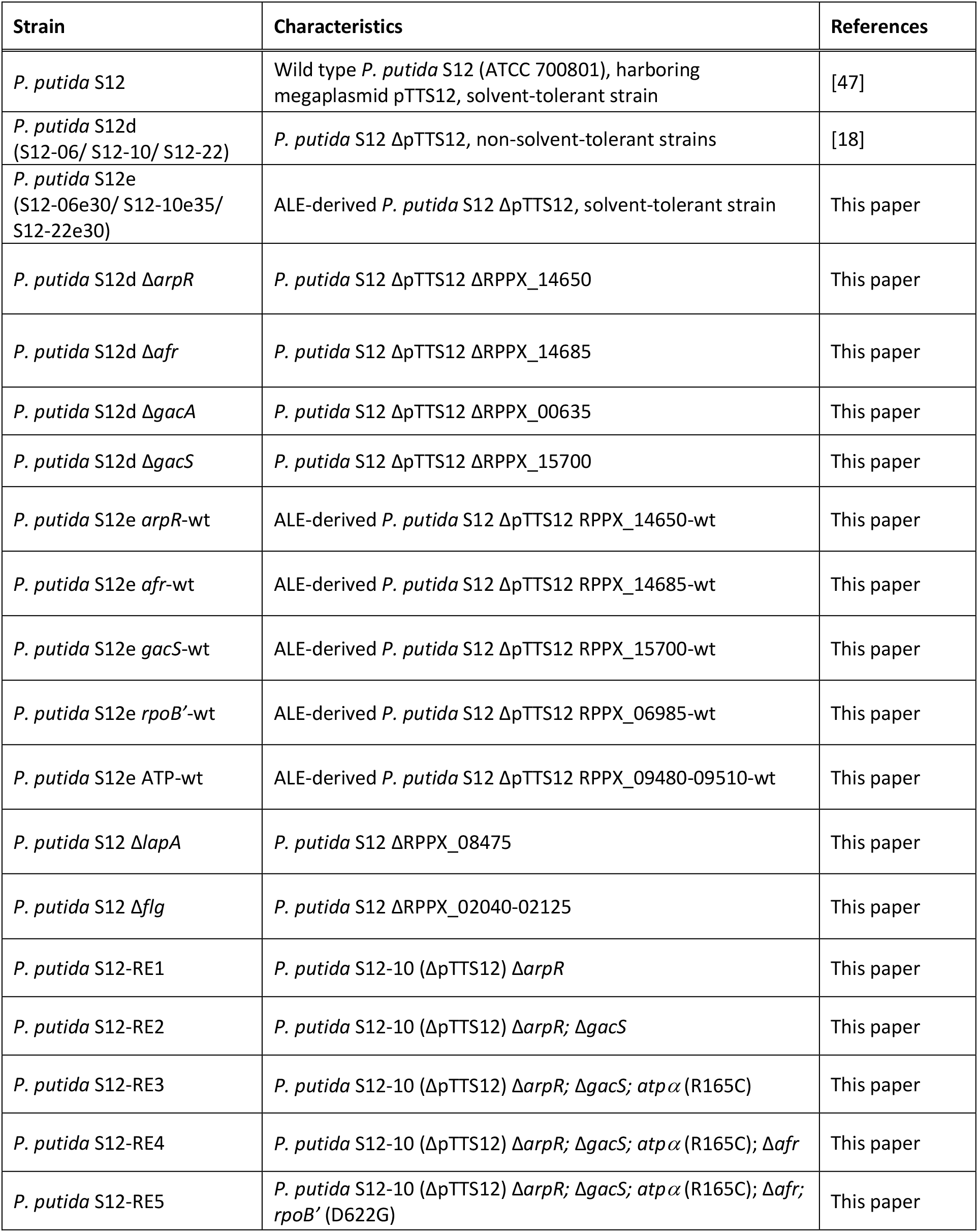
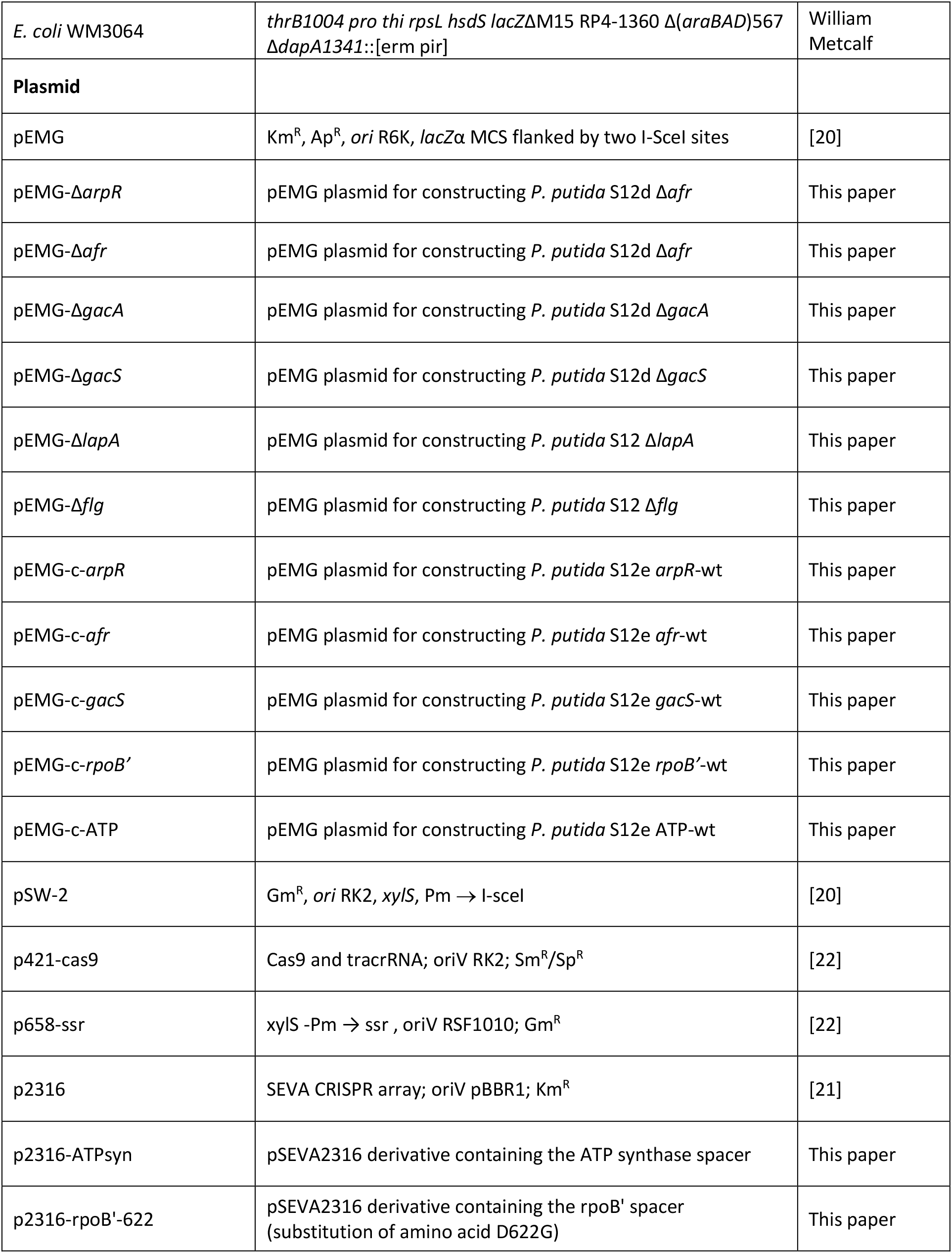
Strains and plasmids used in this paper.

### Adaptive Laboratory Evolution

*P. putida* strains were grown overnight on LB, 30 °C with 200 rpm shaking. Starting cultures were diluted 100 times with LB (starting OD_600_ ± 0.05) and 20 ml of this diluted cultures were placed in Boston bottles. Toluene was added (0.15% v/v) into the cultures and the bottles were immediately closed using Mininert^®^ bottle caps. These cultures were grown on 30 °C with 200 rpm shaking for approximately 24-48 hours to allow the strains to reach exponential phase. The toluene-adapted cultures were then diluted 100 times with LB and grown overnight on 30 °C with 200 rpm shaking. Stocks were made from this LB culture and the cycle of toluene adaptation were continued with higher toluene concentration (0.2% v/v). This cycle was repeated up to 8 cycles with the addition of 0.5% v/v toluene as shown on Figure 1A.

**Figure 1.**
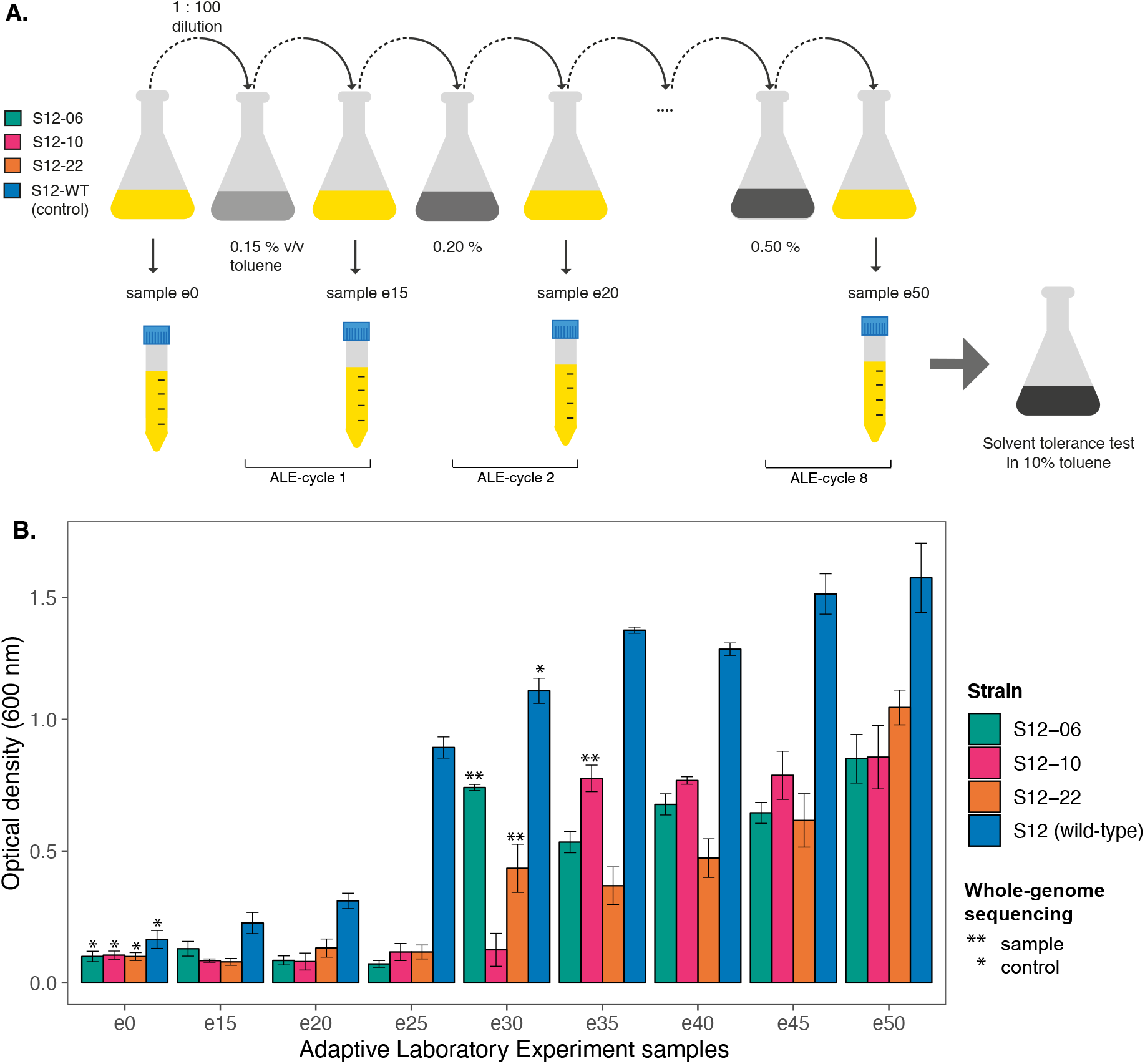
Adaptive laboratory evolution (ALE) experiment of the plasmid-cured *P. putida* S12 to increasing concentration of toluene. A. Experimental design of ALE. ALE was performed on three plasmid-cured *P. putida* S12 strains (S12-06, S12-10, and S12-11). In the ALE experiment, LB media (yellow) was used as the growth media with the addition of increasing toluene concentration 0.05% v/v every cycle (grey). B. Plasmid-cured *P. putida* S12 regained the ability to grow on high toluene concentration. The solvent-tolerance phenotype of ALE-derived strains was tested by observing strain growth on LB with 10% v/v toluene within 48 hours. The asterisks (*) and double asterisk (**) indicate the control and sample strains that were taken for whole genome sequencing. This experiment was performed with three biological replicates and error bars indicate standard deviation.

### PCR and cloning methods

PCR reactions were performed using Phusion polymerase (Thermo Fisher) according to the manufacturer’s manual. Oligos used in this paper (Table 2) were procured from Sigma-Aldrich. PCR products were analysed by gel electrophoresis on 1 % (w/v) TBE agarose containing 5 μg mL^−1^ ethidium bromide (110V, 0.5x TBE running buffer).

**Table 2.**
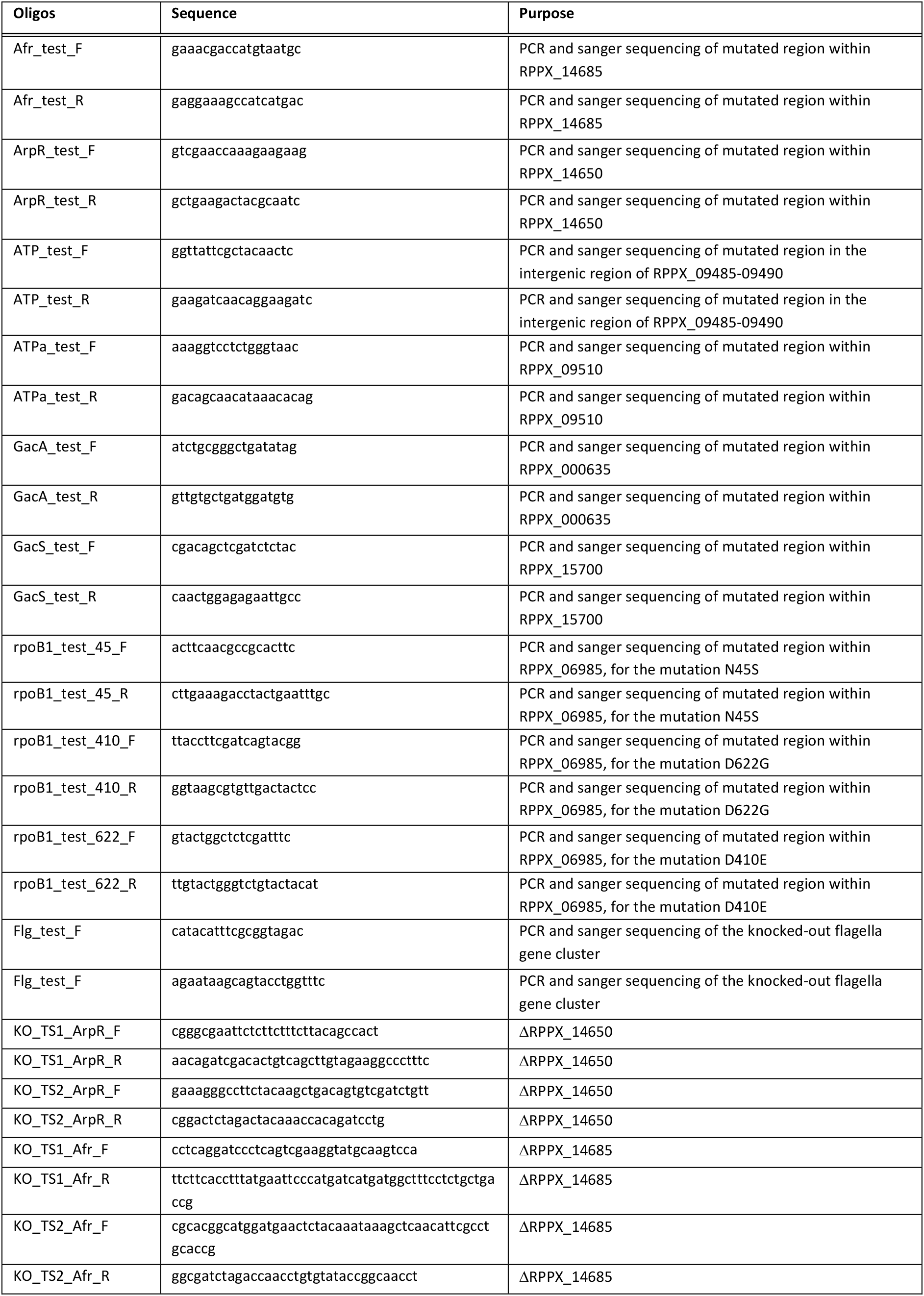
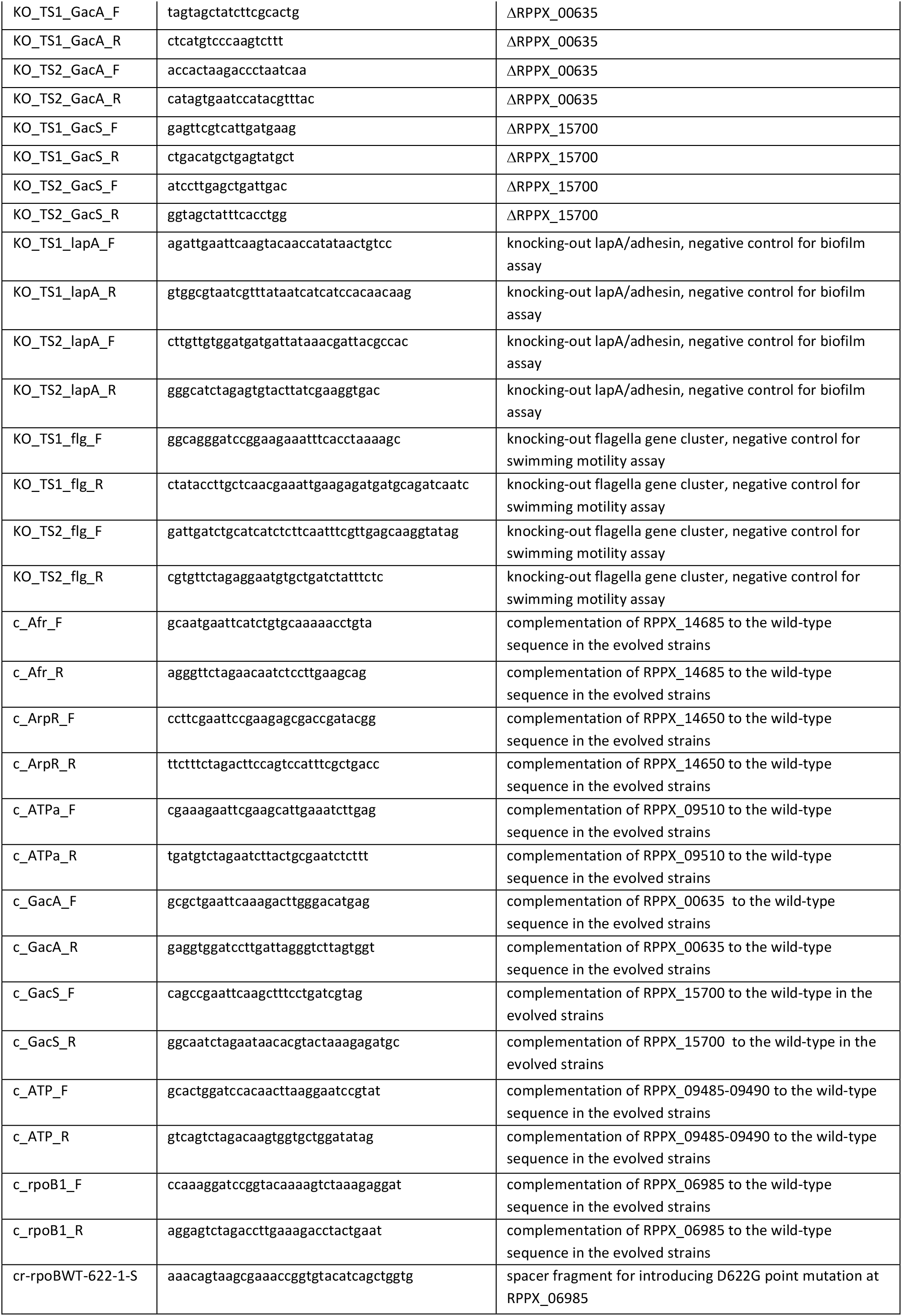
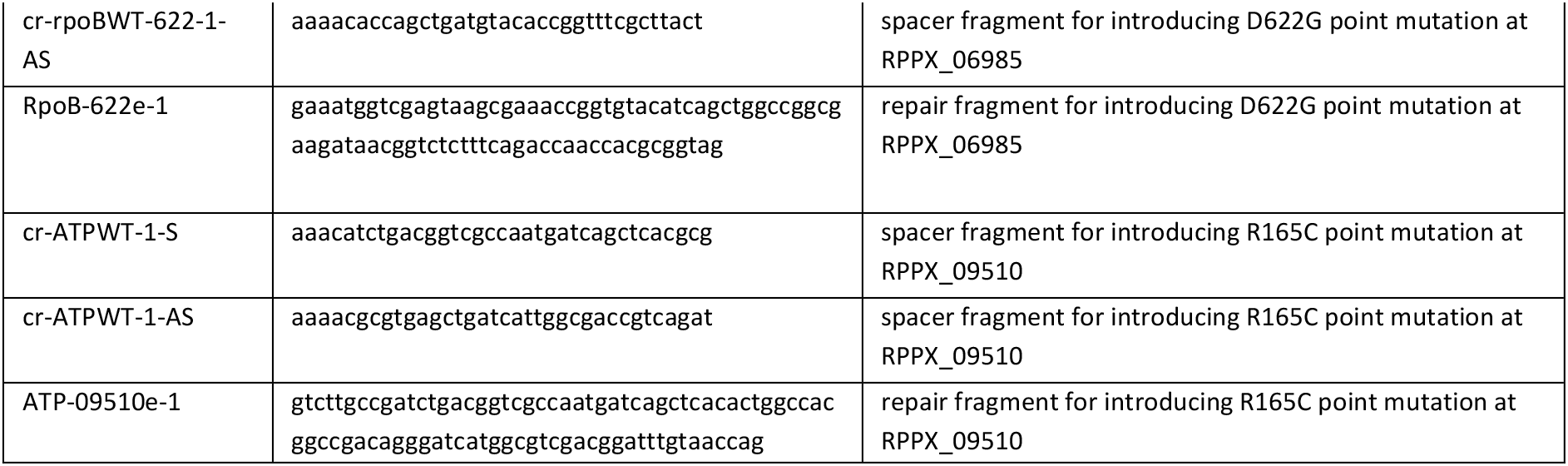
Oligos used in this study.

Deletion of *arpR*, *gacA*, *gacS*, and *afr* genes and restoration of mutations on ALE-derived strains were performed using homologous recombination between free-ended DNA sequences that are generated by cleavage on unique I-SceI sites [20]. Two homologous recombination fragments (TS-1 and TS-2) were obtained by performing PCR using oligos listed in Table 2. Reverse engineering of the point mutations at ATP synthase subunit α (RPPX_09510) and RNA polymerase subunit β’ (RPPX_06985) loci were performed using CRISPR-cas9 enhanced single stranded DNA (ssDNA) recombineering method [21, 22]. Spacers and ssDNA repair fragments were created using oligos listed in Table 2. All of the obtained plasmid constructs, deletion, and restoration of the selected genes were verified by Sanger sequencing (Macrogen BV., Amsterdam).

### Whole-genome sequencing of plasmid-cured and ALE-derived strains

For whole genome sequencing, DNA was extracted by phenol-chloroform extraction followed by column clean-up using NucleoSpin^®^ DNA Plant II kit (Macherey-Nagel). Clustering and DNA sequencing of wild-type, plasmid-cured, and ALE-derived *P. putida* S12 strains were performed using Illumina cBot and HiSeq 4000 (GenomeScan BV, The Netherlands). Image analysis, base calling, and quality check were performed with the Illumina data analysis pipeline RTA v.2.7.7 and Bcl2fastq v.2.17. Raw-data reads were assembled according to the existing complete genome sequence (Accession no. CP009974 and CP009975) in Geneious software [16]. The raw sequence data have been submitted to the SRA database under accession number PRJNA602416.

### RNA sequencing of plasmid-cured and ALE-derived strains

Wild-type, plasmid-cured, and ALE-derived *P. putida* S12 cultures were grown from overnight culture (100 times diluted) on 20 ml LB for 2 hours (30 °C, 200rpm) with and without the addition of 0.1% v/v toluene to bacterial cell cultures. RNA was extracted using TRIzol reagent (Invitrogen) according to the manufacturer’s manual. The obtained RNA samples were cleaned-up using NucleoSpin^®^ RNA Plant and Fungi kit (Macherey-Nagel). RNA libraries were prepared for sequencing using standard Illumina protocols and paired-end sequence reads were generated using the Illumina MiSeq system (BaseClear BV, The Netherlands). Initial quality assessment was based on data passing the Illumina Chastity filtering. Subsequently, reads containing PhiX control signal were removed using an in-house filtering protocol. In addition, reads containing (partial) adapters were clipped (up to a minimum read length of 50 bp). The second quality assessment was based on the remaining reads using the FASTQC quality control tool version 0.11.5. Tophat2 version 2.1.1 aligned RNA-seq reads to a reference genome (Acc. No. CP009974 and CP009975) using the ultra-high-throughput short read aligner Bowtie version 2.2.6 [23, 24]. Cufflink was used to test for differential expression and regulation in RNA-Seq samples. Cuffdiff then estimated the relative abundances of these transcripts. Datasets generated from RNA-seq experiment have been submitted to the GEO database under accession number GSE144045.

### Microtiter dish biofilm formation assay

To quantify biofilm formation, crystal-violet based assay on 96-well plate was performed as described by O’Toole [25]. Overnight cultures of *P. putida* S12 (100 times diluted) were grown in flat bottomed 96-well microtiter plate with 100 μl LB media (30 °C) for 6 hours without shaking. After 6 hours, OD_600_ was measured using Tecan Spark^®^ 10M (Tecan) to ensure the growth of *P. putida* S12. Liquid cultures were removed from 96-well microtiter plate and followed by two-times washing with water. Crystal violet solution (0.1% v/v) 125 μl was added to each well followed by 10-15 minutes of incubation. After incubation, crystal violet solution was removed and the wells were washed with water to remove the excess crystal violet. Microtiter plate was then turned upside-down and dried. Acetic acid solution (30% v/v) 125 μl were added to solubilized biofilm stained crystal violet and incubated for 10-15 minutes. The absorbance at 550 nm was measured to represent biofilm formation using Tecan Spark^®^ 10M (Tecan) and acetic acid solution as blank.

### Swimming motility assay

As starting culture, *P. putida* S12 strains were streaked and grown on LB agar overnight (30 °C). Single colonies were picked and stab-inoculated on to low viscosity LB agar (0.3% w/v agar). This agar was incubated cap-side up for 24 hours at 30 °C. Radial growth of *P. putida* S12 on low-viscosity agar was measured with three replicates to represent swimming motility.

## Result

### Plasmid-cured *Pseudomonas putida* S12 can regain the ability to tolerate high-concentration *toluene*

To investigate the intrinsic solvent-tolerance of *P. putida* S12, we performed adaptive laboratory evolution (ALE) experiment on plasmid-cured *P. putida* S12. Three biological replicates of plasmid-cured *P. putida* S12 (strain S12-06, S12-10, and S12-22) and a wild-type *P. putida* S12 as control were set up to grow on lysogeny broth (LB) media with the addition of 0.15% v/v toluene; the initial maximum concentration that can be tolerated by plasmid-cured *P. putida* S12 (Figure 1A). At stationary phase (typically after 24-48 hours), these cultures were transferred (1:100 dilution) to grow overnight on fresh LB media. Overnight LB media cultures were transferred into LB media containing toluene 0.20% (increase of 0.05% toluene) to continue with the next ALE cycle. While plasmid-cured *P. putida* S12 was unable to grow on LB with 0.20% v/v toluene directly, after adaptation to LB with 0.15% v/v toluene these cultures are able to grow on LB with 0.20% v/v toluene. We repeated this growth cycle with increasing concentration every cycle until plasmid-cured *P. putida* S12 strains were able to grow on LB with 0.50% v/v toluene (Figure 1A). All samples from every ALE-cycle were collected and tested for their ability to survive 10% v/v toluene on LB for 48 hours. This concentration was chosen to represent a high toluene concentration which creates a distinct second phase layer in the culture medium.

The plasmid-cured *P. putida* S12 showed improved tolerance in 10% toluene (Figure 1B). After the adaptation with a moderate toluene concentration (0.30-0.35%), plasmid-cured *P. putida* S12 showed a significant increase in their ability to withstand and sustain growth in 10% toluene. ALE-derived strains S12-06e30, S12-10e35, and S12-22e30 were able to grow on LB media with 10% v/v toluene, reaching final OD_600_ of 0.741 ± 0.02, 0.776 ± 0.08, and 0.434 ± 0.158 respectively after 48 hours. These three samples were taken for whole genome sequencing to map the occurring mutations important for the solvent-tolerance phenotype. Wild-type *P. putida* S12, S12e30, and the initial plasmid-cured *P. putida* S12 strains were sequenced as controls.

### Common mutations were identified in solvent-tolerant strains obtained from ALE

We performed whole genome sequencing of ALE-derived strains S12-06e30, S12-10e35, and S12-22e30 to map the occurring mutations that may lead to increased solvent tolerance in the evolved strains. We identified 32, 77, and 5 mutations (SNPs, insertions/deletions, and mobile element ISS12 insertion) respectively, in S12-06e30, S12-10e35, and S12-22e30 (Figure 2A). Among these mutations, four common mutated loci were identified in all strains. These mutations occurred in an AraC-family transcriptional regulator Afr (RPPX_14685), in a RND efflux pump regulator ArpR (RPPX_14650), in RNA polymerase subunit β’ *rpoB*’ (RPPX_06985), and in the intergenic regions and subunits of ATP synthase (RPPX_09480-09510) (Figure 2A and Table S1). Six mutated loci were shared only between S12-06e30 and S12-10e35. Among these six loci, indels occurred within *gacS* locus (RPPX_15700) in S12-06e30 and S12-10e35 while S12-22e30 had a unique SNP within *gacA* locus (RPPX_00635). In *Pseudomonas*, GacS and GacA proteins are known to constitute a two-component regulatory system which regulates biofilm formation, cell motility, and secondary metabolism [26].

**Figure 2.**
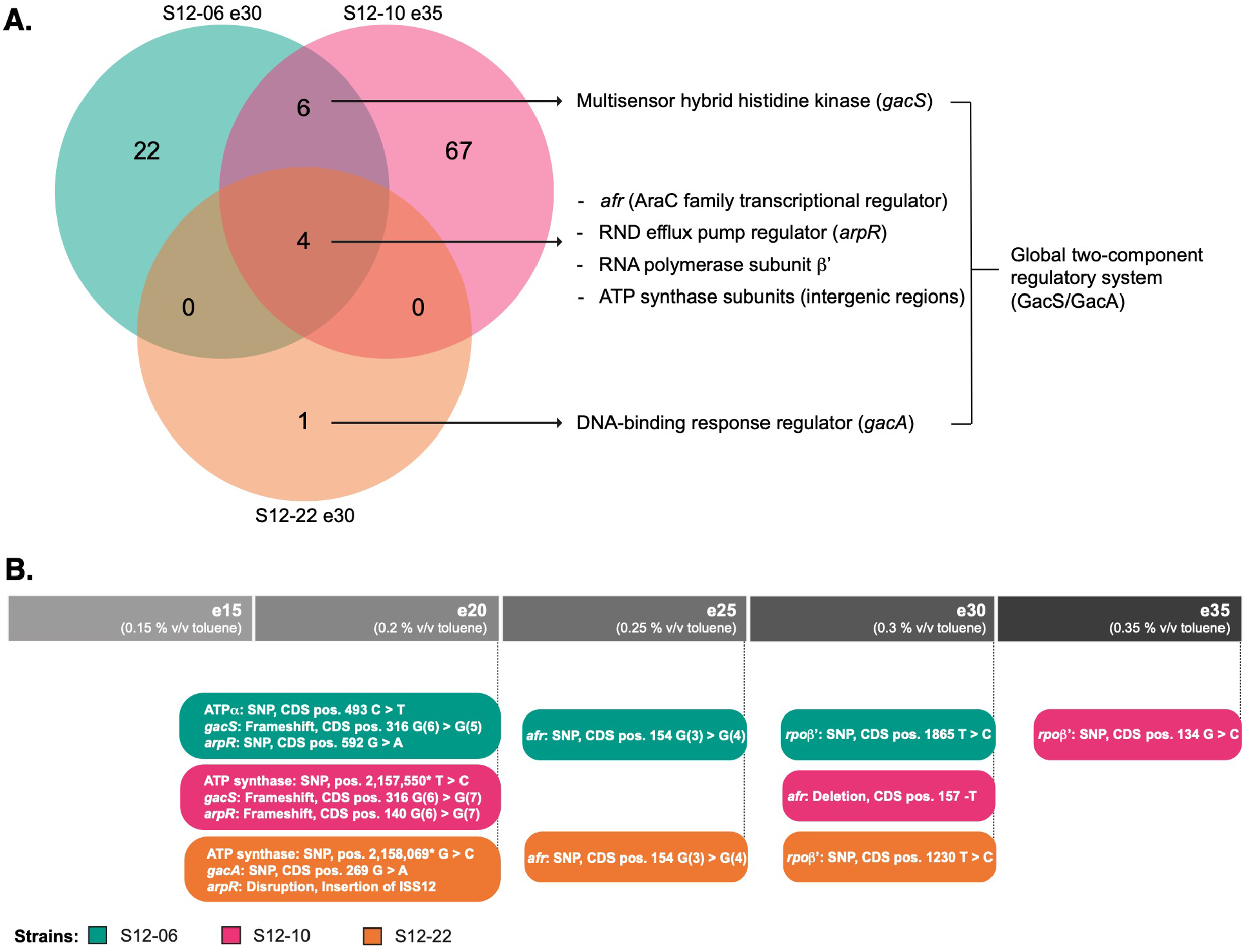
Common mutated loci were identified in the ALE-derived P. putida S12 strains. A. Venn diagram of mutated loci in the ALE-derived *P. putida* S12 strains. The colors indicate the three ALE-derived strains. Common mutated loci were identified among ALE-derived strains on an AraC family transcriptional regulator (Afr), RND efflux pump regulator (ArpR), RNA polymerase subunit β’, intergenic region of F0F1 ATP synthase subunits, and global two-component regulatory system GacS/GacA. B. ALE-derived *P. putida* S12 strains accumulated key mutations in a stepwise manner. ALE-derived strains were probed for key mutations accumulation with PCR and Sanger sequencing. The colors indicate the three ALE-derived strains and the grey bar indicate the ALE cycles which the strains were originating from. The position of the occurring SNPs or indels are indicated as CDS position of each mutated loci, except for the SNPs within the intergenic regions of ATP synthase which positions are indicated relative to the chromosome sequence.

Since the ALE-derived strains showed a sudden increase in their ability to tolerate high toluene concentrations, we investigated the order of accumulation of key mutations in ALE-derived strains. Key mutations accumulated in a stepwise manner rather than emerging simultaneously in one cycle (Figure 2B). In the second ALE-cycle, three key mutations occurred under the exposure to 0.20% v/v toluene in all strains (S12-06e20, S12-10e20, and S12-22e20). The first accumulated mutations occurred on the intergenic regions between ATP synthase subunits, *gacS*/*gacA* loci, and *arpR* locus. In the subsequent cycle, S12-06e25, S12-10e30, and S12-22e25 accumulated additional key mutations in the *afr* locus. The final key mutations *on rpoB’* locus were accumulated by strain S12-06e30, S12-10e35, and S12-22e30, in which the sudden increase of solvent tolerance were observed.

### Contribution of key mutations to increased solvent-tolerance of ALE-derived strains

To study the contribution and impact of each mutated locus, single knock-out strains of *arpR* (RPPX_14650), *afr* (RPPX_14685), *gacA* (RPPX_00635), and *gacS* (RPPX_15700) were created in plasmid-cured *P. putida* S12. In the ALE-derived strains, the acquired mutations (indels and mobile element insertion) in *arpR* (RPPX_14650), *afr* (RPPX_14685), and *gacS* (RPPX_15700) caused truncation of the encoded protein, while the SNP in *gacA* (RPPX_00635) caused an amino acid residue change (P90L) (Figure 2B). The SNPs acquired in ATP synthase and in RNA polymerase subunit β’ loci were not addressed with this single knock-out approach since knocking out these genes would have deleterious effects. Solvent-tolerance analysis of the single knock-out strains indicated that deletion of each of those genes improved the growth of plasmid-cured *P. putida* S12 strains on LB with 0.15% v/v toluene (Figure 3A). However, single knock-out of these genes did not enable plasmid-cured *P. putida* S12 strains to grow on higher toluene concentration than 0.15% v/v.

**Figure 3.**
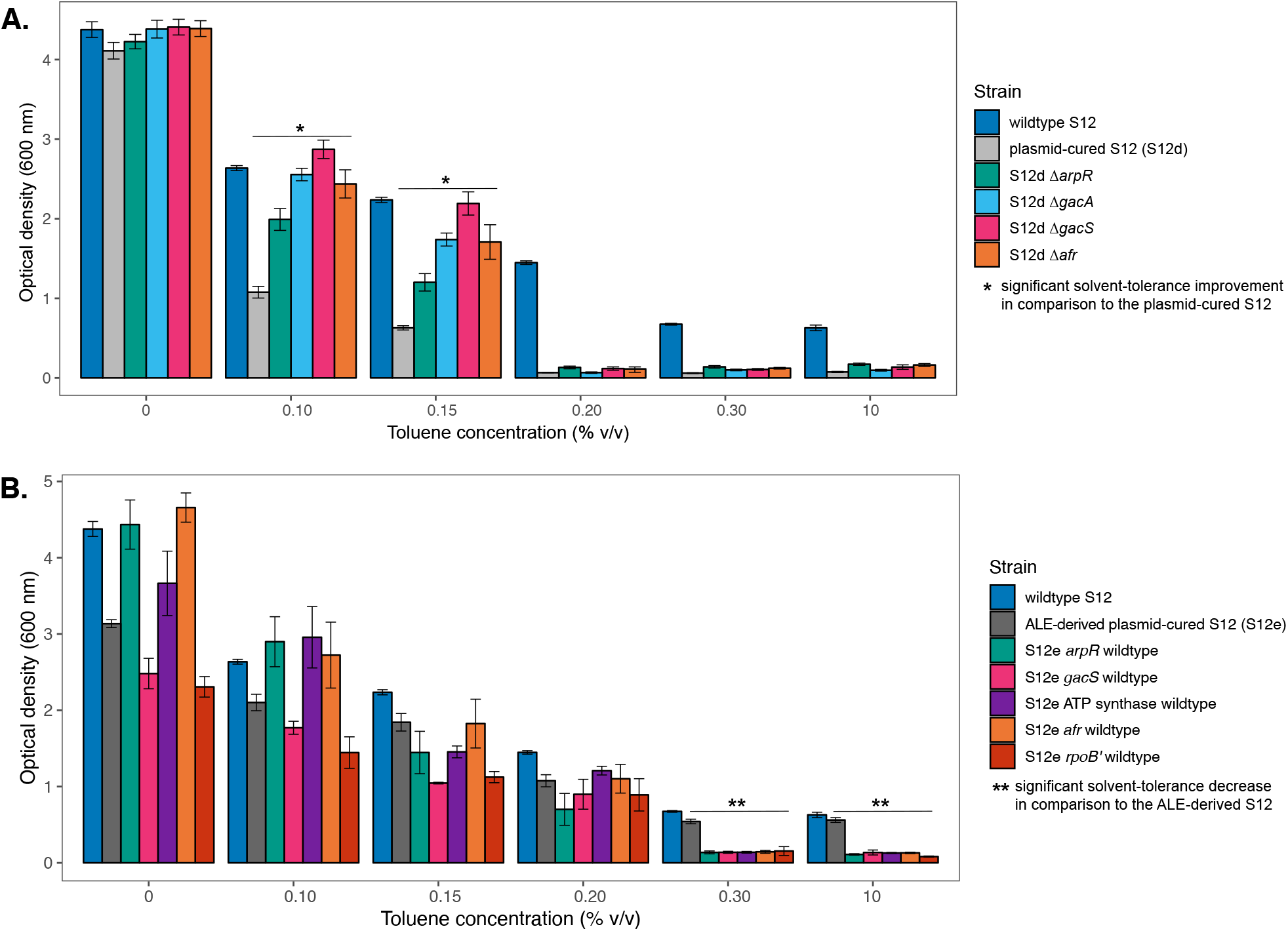
Accumulated key mutations contributed to solvent-tolerance phenotype of ALE-derived P. putida S12 strains. A. Single-knockout of common mutated loci in the plasmid-cured *P. putida* S12 improved strain growth on LB with low toluene concentration (0.1-0.15% v/v). The colors indicate the control strains and the plasmid-cured S12 with deleted loci. This experiment was performed with three biological replicates and error bars indicate standard deviation. B. Single-restoration of common mutated loci in the ALE-derived *P. putida* S12 reduced solvent-tolerance phenotype. The colors indicate the control strains and the ALE-derived S12 with restored loci. The restored strains can grow on LB with a maximum of 0.20% v/v toluene. This experiment was performed with three biological replicates and error bars indicate standard deviation.

Individual restoration of common mutated loci in the ALE-derived strains to their wild-type sequence caused these strains to lose the ability to withstand the presence of moderate and high toluene concentration (0.30% and 10% v/v toluene) (Figure 3B). These strains can sustain growth on LB with maximum 0.20% v/v toluene. Therefore, we concluded that each of the common mutated loci is important for the solvent-tolerance phenotype in ALE-derived strains.

### Reverse engineering of key mutations on plasmid-cured S12 successfully restore solvent-tolerance

To confirm the important contribution of key mutations, we introduced these mutations on a plasmid-cured S12 strain and analyse the growth parameters of the resulting strains in the presence and absence of toluene (Figure 4 and Table 3). Strain S12-10 was chosen to represent the plasmid-cured S12 in this experiment. A knock-out of *arpR* (RPPX_14650) locus was performed resulting in the first reverse engineering strain (RE1). Second knock-out mutation at *gacS* (RPPX_15700) locus was performed on strain RE1, resulting in the strain RE2. It is interesting to note that strain RE2 exhibit significantly better growth parameters in LB and minimal media compared to its parent strains RE1 and S12-10 (Table 3). The third mutation was introduced at the ATP synthase subunit alpha (RPPX_09510) to strain RE2, resulting in the strain RE3. This mutation caused an amino acid substitution from arginine to cysteine at position 165 (R165C), mimicking the mutation found in the strain S12-06e30. Indeed, the introduction of this mutation caused a severe reduction of growth parameters in LB and minimal media (Table 3). Fourth knock-out mutation at *afr* (RPPX_14685) locus was performed on strain RE3, resulting in the strain RE4. Finally, a point mutation was introduced to strain RE4 at *rpoB’* (RPPX_06985) locus to construct strain RE5 which caused an amino acid substitution from aspartic acid to glycine at position 622 (D622G) mimicking the mutation found in the strain S12-06e30.

**Figure 4.**
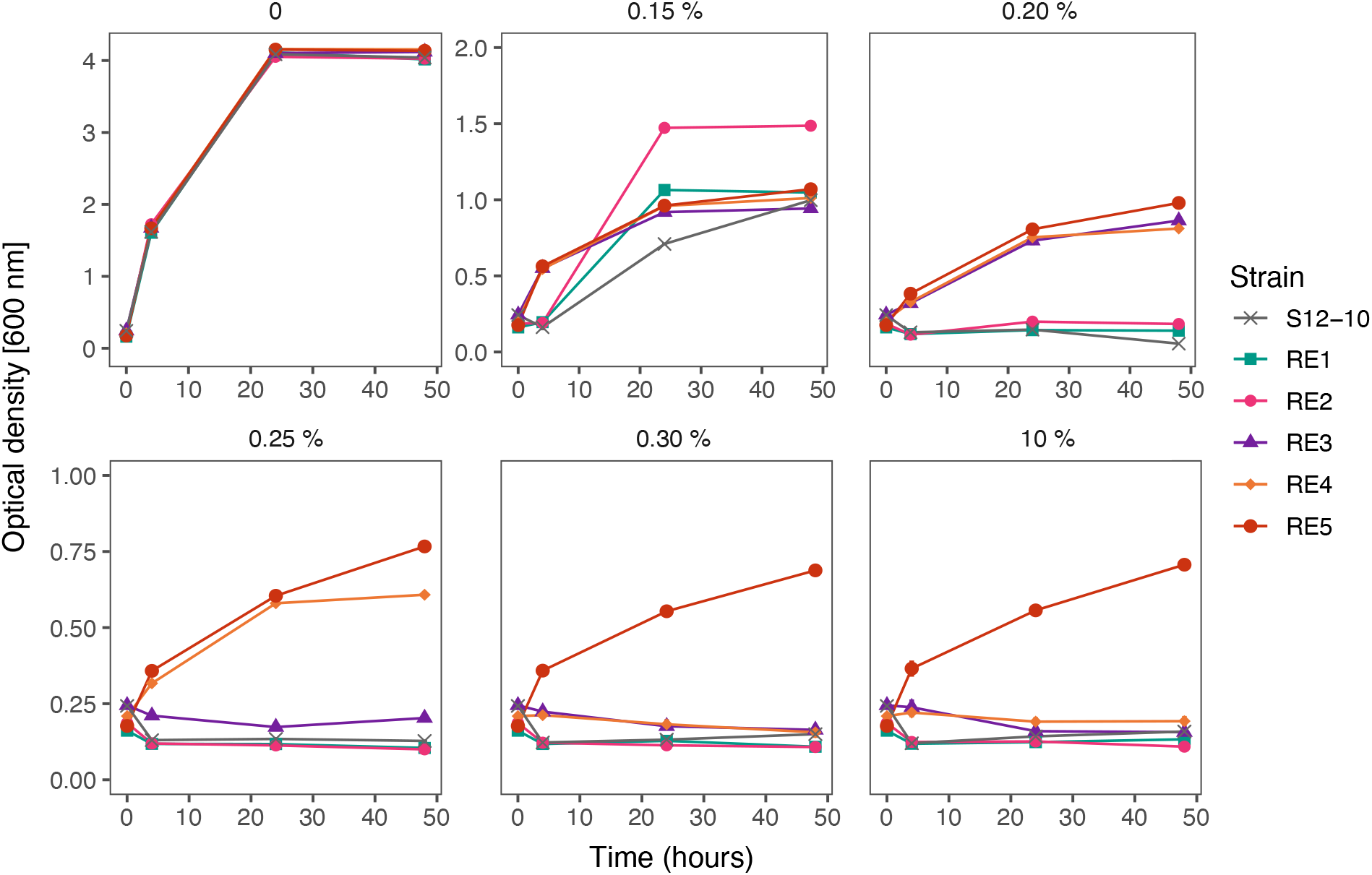
Reverse engineering of the key mutations found in ALE-derived strains. Reverse engineering of the key mutations found in the ALE-derived *P. putida* S12 successfully restores the solvent-tolerance phenotype in the plasmid-cured strain S12-10. The colors indicate the control strain S12-10 and the reverse engineering strains (RE). This experiment was performed with three biological replicates and error bars indicate standard deviation. The y-axis may be different for the presented panels.

**Table 3.**
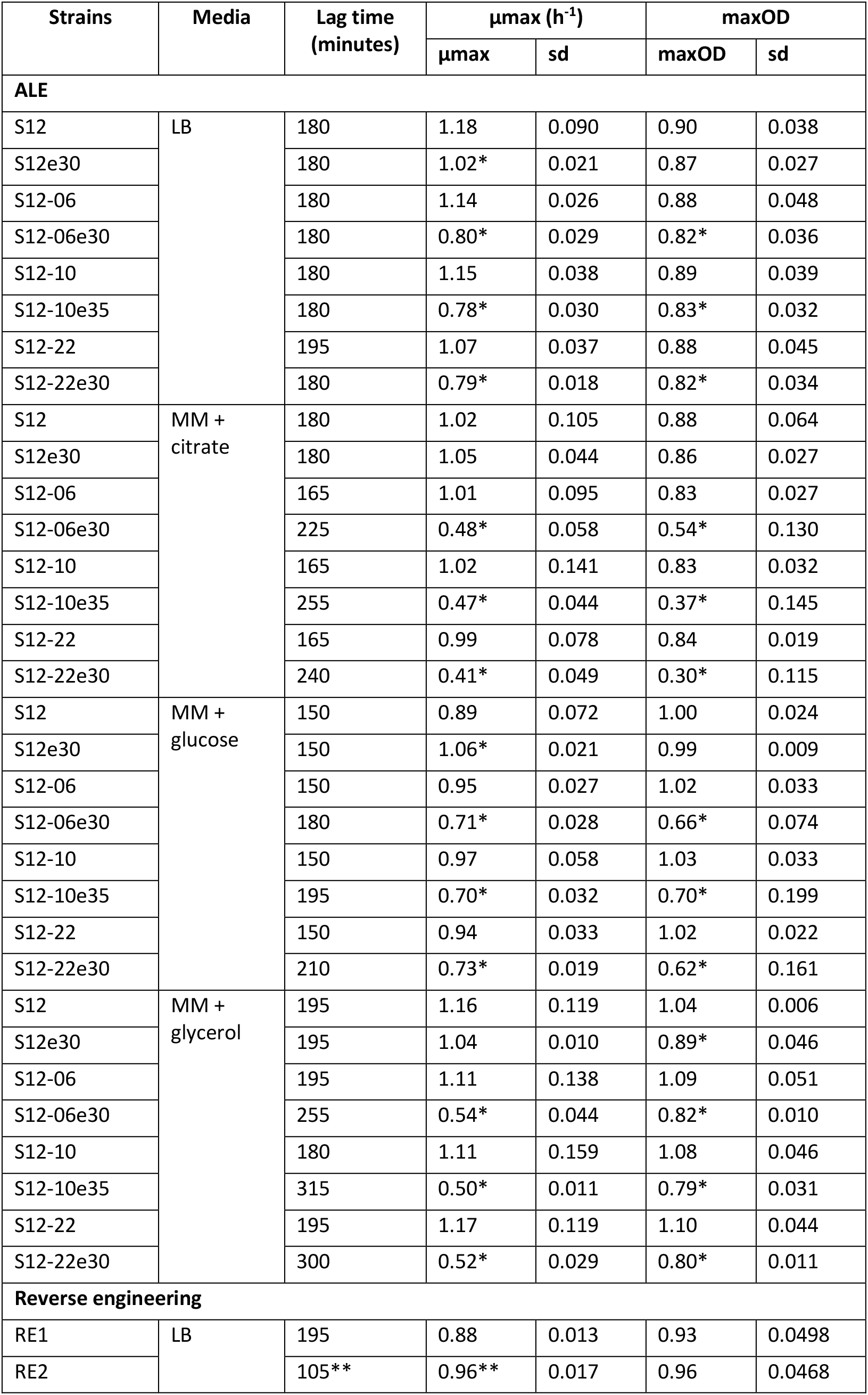
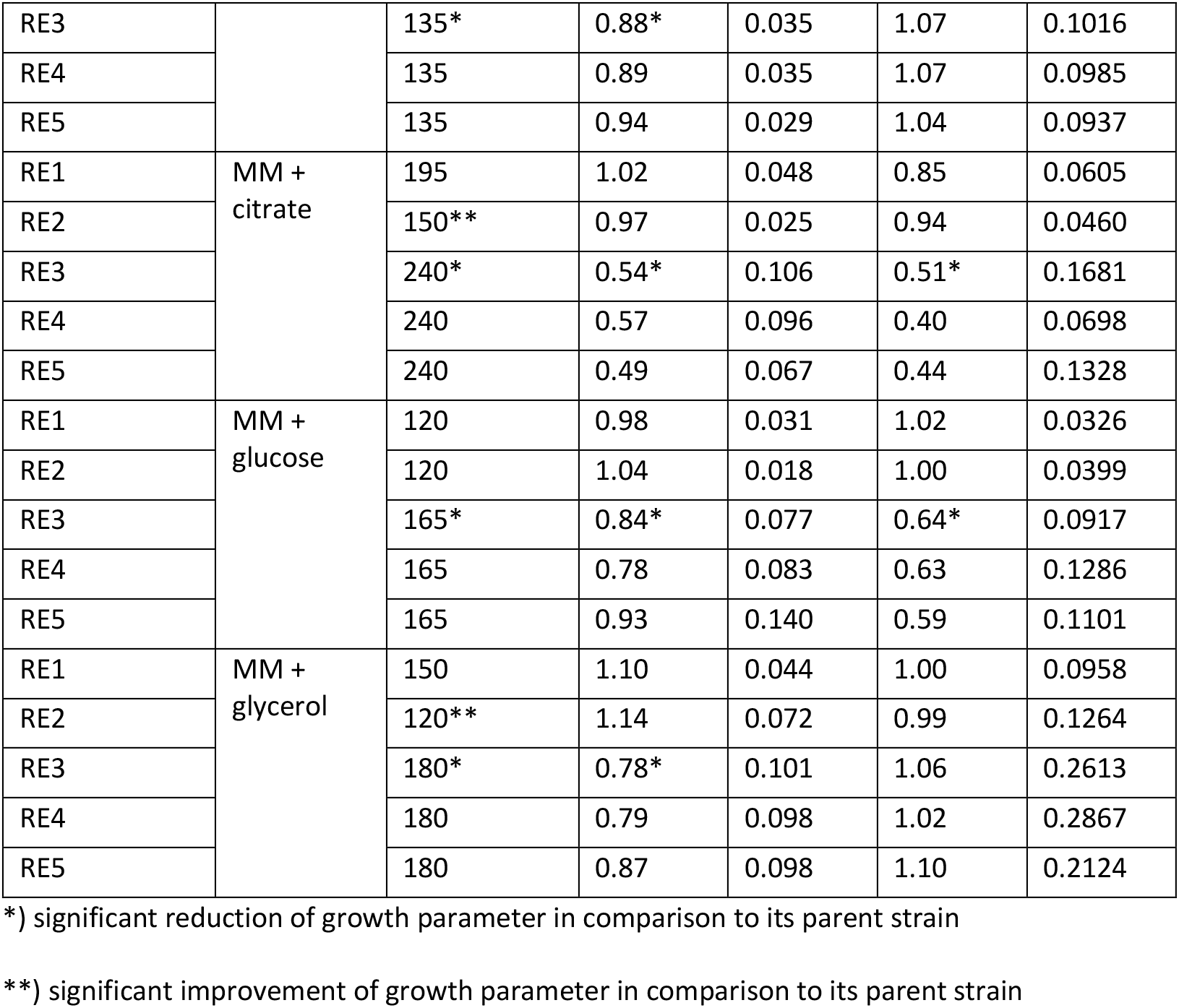
Growth parameters of ALE-derived and reverse engineering strains.

| Strains | Media | Lag time<br>(minutes) | μmax (h<sup>-1</sup>) |  | maxOD |  |
| --- | --- | --- | --- | --- | --- | --- |
|  |  |  | μmax | sd | maxOD | sd |
| ALE |  |  |  |  |  |  |
| S12 | LB | 180 | 1.18 | 0.090 | 0.90 | 0.038 |
| S12e30 |  | 180 | 1.02* | 0.021 | 0.87 | 0.027 |
| S12-06 |  | 180 | 1.14 | 0.026 | 0.88 | 0.048 |
| S12-06e30 |  | 180 | 0.80* | 0.029 | 0.82* | 0.036 |
| S12-10 |  | 180 | 1.15 | 0.038 | 0.89 | 0.039 |
| S12-10e35 |  | 180 | 0.78* | 0.030 | 0.83* | 0.032 |
| S12-22 |  | 195 | 1.07 | 0.037 | 0.88 | 0.045 |
| S12-22e30 |  | 180 | 0.79* | 0.018 | 0.82* | 0.034 |
| S12 | MM +<br>citrate | 180 | 1.02 | 0.105 | 0.88 | 0.064 |
| S12e30 |  | 180 | 1.05 | 0.044 | 0.86 | 0.027 |
| S12-06 |  | 165 | 1.01 | 0.095 | 0.83 | 0.027 |
| S12-06e30 |  | 225 | 0.48* | 0.058 | 0.54* | 0.130 |
| S12-10 |  | 165 | 1.02 | 0.141 | 0.83 | 0.032 |
| S12-10e35 |  | 255 | 0.47* | 0.044 | 0.37* | 0.145 |
| S12-22 |  | 165 | 0.99 | 0.078 | 0.84 | 0.019 |
| S12-22e30 |  | 240 | 0.41* | 0.049 | 0.30* | 0.115 |
| S12 | MM +<br>glucose | 150 | 0.89 | 0.072 | 1.00 | 0.024 |
| S12e30 |  | 150 | 1.06* | 0.021 | 0.99 | 0.009 |
| S12-06 |  | 150 | 0.95 | 0.027 | 1.02 | 0.033 |
| S12-06e30 |  | 180 | 0.71* | 0.028 | 0.66* | 0.074 |
| S12-10 |  | 150 | 0.97 | 0.058 | 1.03 | 0.033 |
| S12-10e35 |  | 195 | 0.70* | 0.032 | 0.70* | 0.199 |
| S12-22 |  | 150 | 0.94 | 0.033 | 1.02 | 0.022 |
| S12-22e30 |  | 210 | 0.73* | 0.019 | 0.62* | 0.161 |
| S12 | MM +<br>glycerol | 195 | 1.16 | 0.119 | 1.04 | 0.006 |
| S12e30 |  | 195 | 1.04 | 0.010 | 0.89* | 0.046 |
| S12-06 |  | 195 | 1.11 | 0.138 | 1.09 | 0.051 |
| S12-06e30 |  | 255 | 0.54* | 0.044 | 0.82* | 0.010 |
| S12-10 |  | 180 | 1.11 | 0.159 | 1.08 | 0.046 |
| S12-10e35 |  | 315 | 0.50* | 0.011 | 0.79* | 0.031 |
| S12-22 |  | 195 | 1.17 | 0.119 | 1.10 | 0.044 |
| S12-22e30 |  | 300 | 0.52* | 0.029 | 0.80* | 0.011 |
| Reverse engineering |  |  |  |  |  |  |
| RE1 | LB | 195 | 0.88 | 0.013 | 0.93 | 0.0498 |
| RE2 |  | 105** | 0.96** | 0.017 | 0.96 | 0.0468 |
| RE3 |  | 135* | 0.88* | 0.035 | 1.07 | 0.1016 |
| RE4 |  | 135 | 0.89 | 0.035 | 1.07 | 0.0985 |
| RE5 |  | 135 | 0.94 | 0.029 | 1.04 | 0.0937 |
| RE1 | MM +<br>citrate | 195 | 1.02 | 0.048 | 0.85 | 0.0605 |
| RE2 |  | 150** | 0.97 | 0.025 | 0.94 | 0.0460 |
| RE3 |  | 240* | 0.54* | 0.106 | 0.51* | 0.1681 |
| RE4 |  | 240 | 0.57 | 0.096 | 0.40 | 0.0698 |
| RE5 |  | 240 | 0.49 | 0.067 | 0.44 | 0.1328 |
| RE1 | MM +<br>glucose | 120 | 0.98 | 0.031 | 1.02 | 0.0326 |
| RE2 |  | 120 | 1.04 | 0.018 | 1.00 | 0.0399 |
| RE3 |  | 165* | 0.84* | 0.077 | 0.64* | 0.0917 |
| RE4 |  | 165 | 0.78 | 0.083 | 0.63 | 0.1286 |
| RE5 |  | 165 | 0.93 | 0.140 | 0.59 | 0.1101 |
| RE1 | MM +<br>glycerol | 150 | 1.10 | 0.044 | 1.00 | 0.0958 |
| RE2 |  | 120** | 1.14 | 0.072 | 0.99 | 0.1264 |
| RE3 |  | 180* | 0.78* | 0.101 | 1.06 | 0.2613 |
| RE4 |  | 180 | 0.79 | 0.098 | 1.02 | 0.2867 |
| RE5 |  | 180 | 0.87 | 0.098 | 1.10 | 0.2124 |

\*) significant reduction of growth parameter in comparison to its parent strain
\*\*) significant improvement of growth parameter in comparison to its parent strain

Reverse engineering strains and its parent, strain S12-10, were tested for their ability to survive and sustain growth in the presence of toluene (Figure 4). Strain S12-10, RE1, and RE2 were only able to withstand and sustain growth in the presence of 0.15% v/v toluene. Nevertheless, strain RE1 and RE2 showed growth improvement in comparison to strain S12-10 (Figure 4, panel 2). Strain RE3 and RE4 were able to withstand and sustain growth in a slightly higher toluene concentration, 0.20% and 0.25% v/v toluene respectively. Finally, strain RE5 were able to sustain growth in the presence of high toluene concentration (10% v/v toluene). These results confirm that the key mutations are indeed important for the restoration of solvent tolerance in plasmid-cured S12.

### Restoration of solvent-tolerance involved a constitutive downregulation of energy consuming activities in ALE-derived strains

Global transcriptional analysis (RNA sequencing) was performed to probe the response of ALE-derived *P. putida* S12 in comparison with wild-type and plasmid-cured *P. putida* S12 in the presence of toluene (LB with 0.1% v/v toluene). As a response to toluene addition, ALE-derived strains showed differential expression only of 14 loci. This response was in stark contrast to the wild-type S12 and plasmid-cured S12 which differentially expressed more than 500 loci as a response to toluene addition (Table S2). Comparisons of gene expression between ALE-derived strains with plasmid-cured and wild-type *P. putida* S12 growing on LB in the absence of toluene indicated that the mutations which occurred in the ALE-derived strains caused constitutive differential expression of ± 900 genes which play a role in restoring solvent tolerance (Table S3).

Constitutive differentially expressed genes in ALE-derived strains in comparison to parental plasmid-cured *P.putida* S12 were classified based of COG categorization. Several classes of genes were downregulated in ALE-derived strains compared to plasmid-cured S12, for example, genes constituting cell motility, intracellular trafficking and secretion, and defence mechanism functions (Figure 5A). In general, ALE-derived strains appeared to constitutively shut-down energy consuming activities, such as flagella biosynthesis, F0F1 ATP synthase, and membrane transport proteins which are energized through proton (H^+^) influx. Additionally, genes related to biofilm formation were constitutively down-regulated. Here, we focused on several classes of genes that were differentially expressed in ALE-derived strains compared to its parental strain.

**Figure 5.**
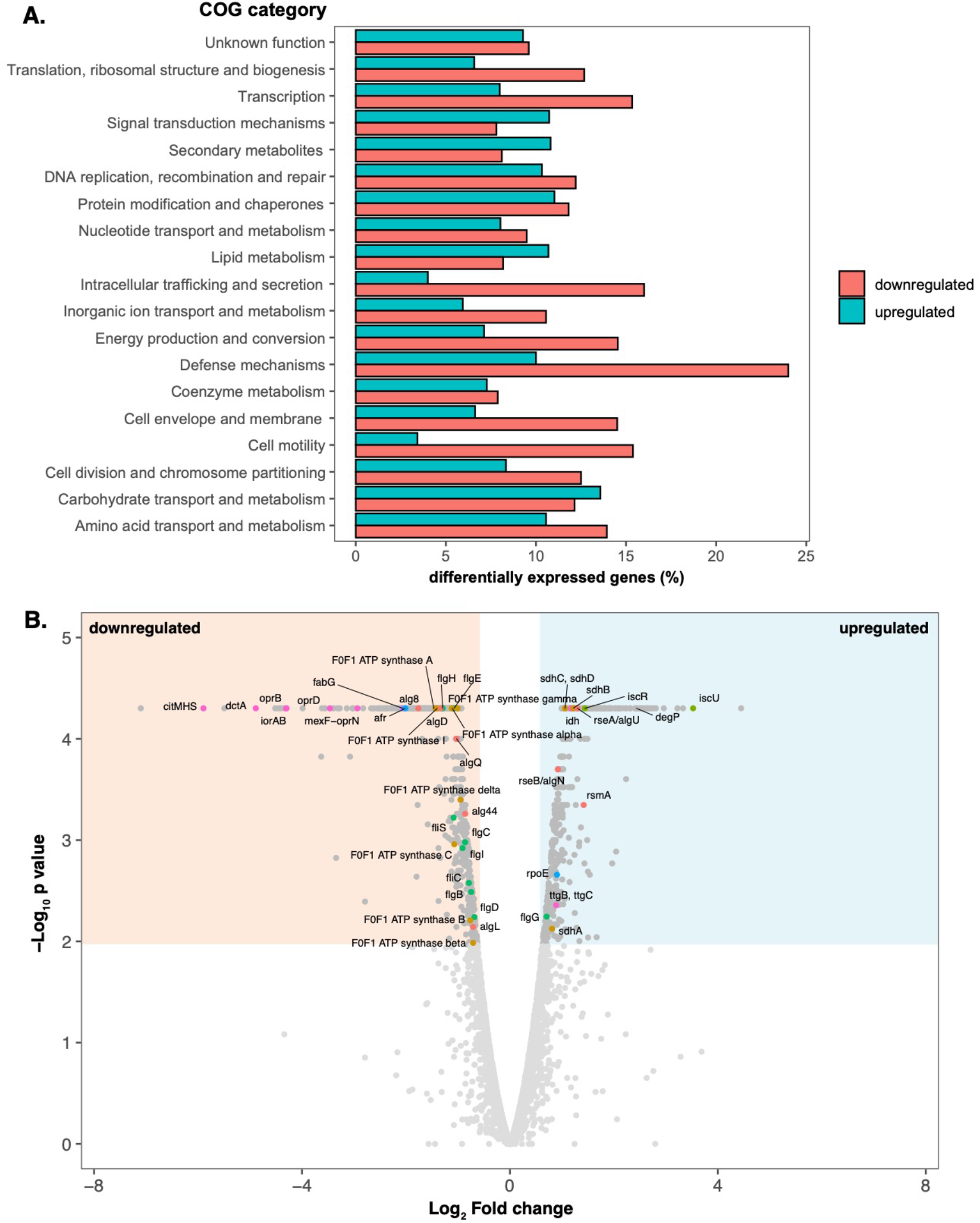
Visualization of differentially expressed genes on ALE-derived P. putida S12 strains in comparison to its parental strain growing on LB. A. COG classification of differential gene expression in ALE-derived strains in comparison to plasmid-cured *P. putida* S12. COG classification was performed using eggNOG 5.0 mapper (http://eggnogdb.embl.de/#/app/emapper). The percentages of up-regulated genes in each class are represented by the blue bar and the percentages of down-regulated genes are represented by the red bar. B. Volcano plot of differential gene expression in ALE-derived strains in comparison to the plasmid-cured *P. putida* S12 growing on LB. Blue area indicates the significantly upregulated genes and beige area indicates the significantly downregulated genes (cut-off: Log2 Fold Change ≥ 1 for up-regulated genes or ≤ 1 for down-regulated genes and p-value ≤ 0.01). The colored dots represent the significantly up-/down-regulated genes discussed in this paper. The colors correspond to the functions of each genes: red represents biofilm and alginate production genes; orange represents the genes involved in oxidative phosphorylation process; light green represents the genes involved in energy production process; green represents flagellar assembly gene clusters; blue represents sigma factor and transcriptional regulator; and magenta represents the genes which constitute membrane transporters.

#### Membrane proteins and efflux pumps

ArpABC efflux pump (RPPX_14635-14640) is a multi-functional RND efflux pump homologous to TtgABC from *P. putida* DOT-T1E. This locus was moderately upregulated in ALE-derived strains (Figure 5B; *ttgB* and *ttgC*). While the upregulation of this pump is a common response to toluene in wild-type *P. putida* S12, ALE-derived strains constitutively upregulate ArpABC by SNPs and mobile element insertion in its negative regulator gene *arpR* (RPPX_14650). Interestingly, almost all of the other RND efflux pumps encoded in the chromosome of *P.putida* S12 were downregulated in ALE-derived strains. In the ALE-derived strains, lacking the pTTS12-encoded SrpABC solvent pump, ArpABC is the only remaining efflux pump that may extrude toluene, albeit with a much lower affinity [27]. Downregulation of other efflux pumps is likely to be important to preserve the required proton motive force.

The genes associated with porin function were downregulated in ALE-derived strains as exemplified by RPPX_10240, RPPX_14820, and RPPX_17640 which encodes OprD porin family proteins. This response was similar to previous proteomics study [28] which noted the downregulation of porins to avoid toluene leakage into the cell through these porins. In addition to porins downregulation, several membrane transport proteins like *dctA* (H^+^/C4-dicarboxylate symporters, RPPX_17630) and *citMHS* (citrate-divalent cation/H^+^ symporter, RPPX_17635) were constitutively downregulated.

#### Energy production and conversion

In ALE-derived strains, F0F1 ATP synthase subunits were constitutively downregulated (Figure 5B). This was in-line with our finding of the SNPs which occurred on the intergenic regions between F0F1 ATP synthase subunits (RPPX_09480-RPPX_09510). F0F1 ATP synthase generates 1 ATP from ADP in bacteria by pumping out 3 H^+^ molecule and thus downregulation of this loci may also contributed to the preservation of proton motive force.

Succinate dehydrogenase (SdhABCD) gene cluster (RPPX_01070-RPPX_01085) were constitutively upregulated in ALE-derived strains. Succinate dehydrogenase is responsible as complex II in oxidative phosphorylation process. Cytochrome C oxidase subunit II (RPPX_08860) and its assembly protein (RPPX_08850), composing complex IV, were also constitutively upregulated in ALE-derived strains. Taken together, these findings underlined the importance of electron transport chain in maintaining proton motive force during solvent-stress.

#### Biofilm formation

In ALE-derived strains, we observed a constitutive upregulation of the *rsmA* locus (RPPX_02245). Upregulation of *rsmA* locus may be caused by the mutations found in *gacS*/*gacA* locus and is known to promote motile lifestyle in *Pseudomonas* [26]. Downregulation of alginate biosynthesis pathway as the main polysaccharide matrix in *Pseudomonas* biofilm was also observed. Alg44 (RPPX_14155), which upon its interaction with c-di-GMP is known to positively regulate alginate production [29], was constitutively downregulated in ALE-derived strains. Other loci which are involved in alginate biosynthesis and export were also down regulated e.g. *algL* (RPPX_14130), *alg8* (RPPX_14160), *algD* (RPPX_14165), and *algE* (RPPX_21545). Taken together, these findings pointed to reduction of biofilm formation capacity in the ALE-derived strains.

To confirm this result, we applied a microtiter dish biofilm formation assay [25] to assess the biofilm formation in ALE-derived strains (Figure 6A). Biofilm formation was indeed clearly lower in ALE-derived strains compared to the wild-type and plasmid-cured *P. putida* S12. This tendency was reversed when the indel mutation in the *gacS* locus was restored to the wild-type sequence. While biofilm may protect bacteria from external stressors, nutrient and oxygen depletion during sessile lifestyle may be disadvantageous in solvent-stress and therefore constitutive downregulation of biofilm-related genes was beneficial in ALE-derived strain.

**Figure 6.**
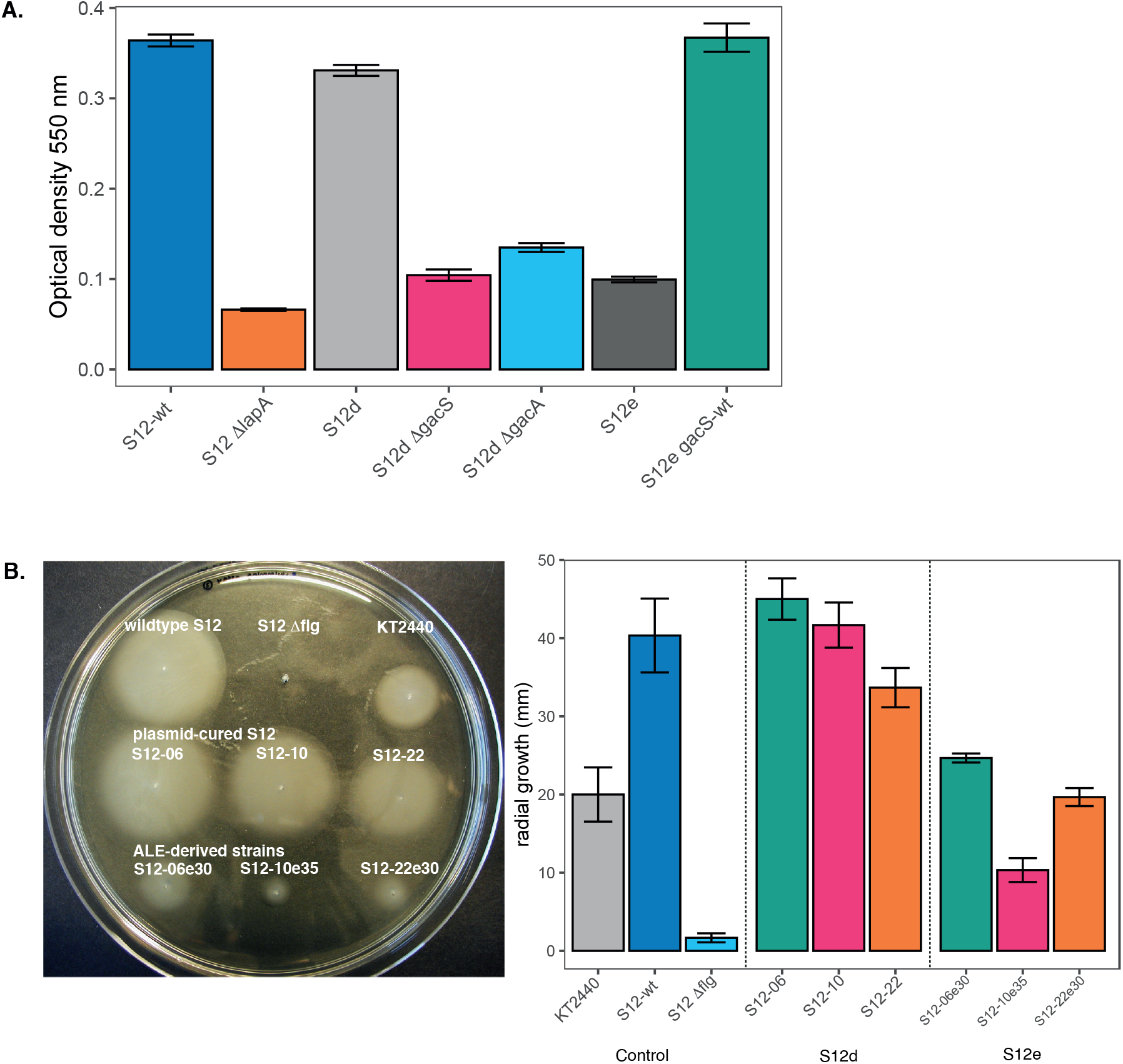
Biofilm formation and cell motility were reduced in ALE-derived P. putida S12 strains. A. Microtiter biofilm formation assay of *P. putida* S12. Plasmid-cured *P. putida* S12 (S12d) ΔgacS and ΔgacA showed similar reduction as ALE-derived *P. putida* S12 (S12e) strains. Restoration of *gacS* locus to wild-type sequence (S12e *gacS*-wt) also restore biofilm formation in ALE-derived *P. putida* S12 (S12e). The measurement of biofilm formation was performed by measuring optical density at 500nm as previously described [25] with ΔlapA (adhesin) taken as negative control. This experiment was performed with three biological replicates and error bars indicate standard deviation. B. Swimming motility assay of *P. putida* S12 in low viscosity agar (LB + 0.3% agar). ALE-derived *P. putida* S12 strains (S12e) showed a reduced radial growth in low viscosity agar indicating lower swimming motility. The Δflg (flagella gene cluster) was taken as a negative control. On the second panel, the bars represent an average of radial growth of at least three biological replicates of each strains and error bars indicate standard deviation.

#### Cell motility

Flagellar biosynthesis loci (RPPX_02045-RPPX_02125) were constitutively downregulated in ALE-derived strains (Figure 5B). Consequently, this may lead to reduced swimming motility in ALE-derived strains. We confirmed this finding by measuring the radial growth of ALE-derived strains in comparison to the wild-type and plasmid-cured *P. putida* S12 on low-viscosity agarose (Figure 6B). Indeed, ALE-derived strains showed a significant reduction of radial growth. Downregulation of flagella may be a strategy of ALE-derived strains to maintain proton motive force and reroute its energy towards extrusion of toluene since both RND efflux pump ArpABC and flagella utilize H^+^ influx as energy source.

#### Chaperones

We also observed the constitutive upregulation of loci RPPX_14680-14875 which encode homologs of sigma factor E (RpoE), anti-sigma factor RseAB, and DegP protein respectively. This cluster is known to orchestrate the expression of chaperone proteins as a stress response to the elevated amount of misfolded proteins in *E.coli* [30]. Additionally, these sigma factors are known to negatively regulate alginate biosynthesis in *P. aeruginosa* [31]. Other chaperone protein, like Hsp20 protein (RPPX_17155) was also constitutively upregulated in ALE-derived strains. Constitutive upregulation of these genes suggested an important role of chaperones in the adaptive response to high toluene concentration.

## Discussion

### Solvent tolerance can be restored in a relatively small number of generations

The single copy megaplasmid pTTS12 plays an essential role in the solvent-tolerance trait of *P. putida* S12. An efficient solvent extrusion pump SrpABC (homologous to TtgGHI), styrene-phenylacetate degradation pathway, and the recently-identified toxin-antitoxin SlvTA are encoded within this megaplasmid [16]. Unlike *P. putida* DOT-T1E, *P. putida* S12 does not encode toluene degradation pathway within its genome and thus, its solvent-tolerance heavily relies on the gene clusters encoded in pTTS12 as mentioned above [32]. However, previous attempts expressing SrpABC in other non-solvent tolerance bacteria like *E. coli* were unsuccessful to incite the same level of solvent-tolerance as with *P. putida*. This may indicate that *P. putida* S12 is intrinsically solvent tolerant to begin with [33, 34]. Hence, in this paper, we further scrutinized this putative intrinsic solvent tolerance in *P. putida* S12 using Adaptive Laboratory Evolution (ALE).

Upon curing of megaplasmid pTTS12, solvent-tolerance of *P. putida* S12 was significantly reduced. After 4-5 adaptation cycles (± 7 generations per growth cycle) to increasing toluene concentrations, solvent-tolerance trait of plasmid-cured *P. putida* S12 could be restored. Relatively short adaptation to alternating cycles of LB in the presence or absence of toluene can restore solvent-tolerance to elevated concentration of toluene due to the stringent selection pressure elicited by this experimental set-up. However, we also observed a severe reduction in growth parameters on the resulting ALE-derived strains grown in the absence of toluene compared to wild-type *P. putida* S12 undergoing the same adaptation cycles to toluene. Several common mutated loci were identified between replicates of ALE-derived *P. putida* S12. Whole genome sequencing revealed SNPs, indels, and an insertion of mobile genetic element ISS12 occurred in a negative regulator of RND efflux pump ArpR, an uncharacterized AraC family transcriptional regulator Afr, RNA polymerase subunit β’, intergenic region of F0F1 ATP synthase subunits and global two-components regulatory system GacS/GacA. Each of these mutations were demonstrated to be essential in restoring solvent-tolerance of the plasmid-cured *P. putida* S12.

### Up-regulation of solvent efflux pump is compensated by down-regulation of other membrane proteins

RNA-sequencing revealed constitutive differential changes of gene expression in ALE-derived strains caused by the observed mutations. Truncation of ArpR caused a moderate upregulation of ArpBC locus, confirming the promiscuous function of ArpABC efflux pump as antibiotic pump and solvent pump as was previously described [35]. However, other RND efflux pumps were generally downregulated in ALE-derived strains. Indeed, it has been described that a combination of different efflux pumps expression can be toxic to bacteria [36]. While there are multifactorial causes of efflux pumps toxicity, including membrane composition changes and insertion machinery overload [36–38], we propose that the cause of efflux pump toxicity may be due to a high demand of proton motive force. In ALE-derived strains, F0F1 ATP synthase subunits, flagella and other H^+^ influx-dependent membrane transporters were severely downregulated following the moderate upregulation of ArpBC locus. Downregulation of F0F1 ATP synthase subunits may contribute to the observed fitness reduction and at the same time, required as a strategy in ALE-derived strains to overcome efflux pump toxicity in supporting the immense effort of solvent extrusion.

### Truncation of putative regulator Afr in ALE-derived strains reduces expression of membrane proteins

Indel mutations were observed in a hitherto uncharacterized AraC-family transcriptional regulator (Afr), causing it to be truncated in the ALE-derived strains. In *P.putida* KT2440, a homolog of Afr encoded by PP1395 (100% identity, 100% coverage) was found to be responsible for a decrease in glycerol uptake [39], while in *P. aeruginosa* PA14, Afr homolog (63% identity, 88% coverage) encoded by PA14_38040 (PA2074 in strain PAO1) was reported to regulate the expression of RND efflux pump MexEF-OprN [40]. Further characterization of Afr is underway. Since the above mentioned homologs of Afr pointed to a function regulation of transporters, truncation/deletion of Afr may contribute to the maintenance of proton motive force in ALE-derived strains.

### Mutations in the gacS/gacA loci as a common strategy for swift phenotypic switching in Pseudomonads

Truncation of GacS protein and a SNP at *gacA* locus in ALE-derived strains resulted in the observed upregulation of its target, *rsmA* locus. Alginate biosynthesis genes, the main polysaccharide constituting Pseudomonas biofilm [29], were constitutively downregulated in ALE-derived strains. Indeed, we observed reduced biofilm formation in ALE-derived strains, which could be reversed when the mutation in *gacS* locus was complemented with wild-type sequence. In *P. aeruginosa*, biofilm dispersion can be triggered by carbon starvation and involves proton motive force dependent step(s) [41]. During solvent-stress, efficient carbon catabolism and energy production is essential for the extrusion of solvent and survival of *P. putida* S12, therefore biofilm formation causing carbon starvation and oxygen depletion will be disadvantageous.

In the reverse engineering strain RE2, the deletion of *gacS* locus resulted in a significant improvement of growth parameters in the presence and absence of toluene. This mutation may have been selected for to compensate for the mutations that severely affect the growth of ALE-derived strains, for example, the mutations at F0F1 ATP synthase loci. Similar observation was also reported in previous studies, e.g. the loss of function mutation at the *gacS*/*gacA* loci increased the fitness of plasmid-carrying bacterial strain [42] and improved growth characteristics and efficient root colonization [43, 44]. The GacS/GacA two-component system may have a pleiotropic effect since this system regulates a large set amount of genes as a response to environmental stimuli. Additionally, *gacA*/*gacS* loci may constitute a commonly mutated loci which have an elevated mutation rate to allow for a swift phenotypic switching in the environmental dynamics [42–44].

### General summary

In summary, ALE presents a powerful combination of mutation selection and construction of beneficial genetic variation in many different genes and regulatory regions in parallel [45, 46], for the restoration of solvent-tolerance in plasmid-cured *P. putida* S12. Through ALE, we gained insight into intrinsically promoting solvent-tolerance of *P. putida* S12. High metabolic flexibility of *P. putida* S12, e.g. ability to maintain proton motive force, indeed proved essential to incite solvent-tolerance with the availability of a solvent extrusion pump. This may very well be under the control of *gacA*/*gacS* loci and may involve the putative regulator Afr. Further characterization of the efficiency of solvent extrusion pumps and their impact and demand on proton motive force is required for the application of solvent-tolerant strains, especially in the bioproduction of high-value chemicals and biofuels.

## Acknowledgements

H. Kusumawardhani was supported by the Indonesia Endowment Fund for Education (LPDP) as scholarship provider from the Ministry of Finance, Indonesia. R. Hosseini was funded by the Dutch National Organization for Scientific Research NWO, through the ERAnet-Industrial Biotechnology program, project ‘Pseudomonas 2.0’. We would like to thank Dr. Esteban Martínez-García and Prof. Victor de Lorenzo for kindly providing us with the materials to perform CRISPR-Cas9 gene editing.

